# H_2_ is a Major Intermediate in *Desulfovibrio vulgaris* Corrosion of Iron

**DOI:** 10.1101/2022.11.15.516606

**Authors:** Trevor L. Woodard, Toshiyuki Ueki, Derek R. Lovley

## Abstract

*Desulfovibrio vulgaris* has been the primary pure culture sulfate reducer for developing microbial corrosion concepts. Multiple mechanisms for how it accepts electrons from Fe^0^ have been proposed. We investigated Fe^0^ oxidation with a mutant of *D. vulgaris* in which hydrogenase genes were deleted. The hydrogenase mutant grew as well as the parental strain with lactate as the electron donor, but unlike the parental strain was not able to grow on H_2_. The parental strain reduced sulfate with Fe^0^ as the sole electron donor, but the hydrogenase mutant did not. H_2_ accumulated over time in Fe^0^ cultures of the hydrogenase mutant and sterile controls, but not in parental strain cultures. Sulfide stimulated H_2_ production in uninoculated controls apparently by both reacting with Fe^0^ to generate H_2_ and facilitating electron transfer from Fe^0^ to H^+^. Parental strain supernatants did not accelerate H_2_ production from Fe^0^, ruling out a role for extracellular hydrogenases. Previously proposed electron transfer between Fe^0^ and *D. vulgaris* via soluble electron shuttles was not evident. The hydrogenase mutant did not reduce sulfate in the presence of Fe^0^ and either riboflavin or anthraquinone-2,6-disulfonate and these potential electron shuttles did not stimulate parental strain sulfate reduction with Fe^0^ as the electron donor. The results demonstrate that *D. vulgaris* primarily accepts electrons from Fe^0^ via H_2_ as an intermediary electron carrier. These findings clarify the interpretation of previous *D. vulgaris* corrosion studies and suggest that H_2_-mediated electron transfer is an important mechanism for iron corrosion under sulfate-reducing conditions.

**Importance:** Microbial corrosion of iron in the presence of sulfate-reducing microorganisms is economically significant. There is substantial debate over how microbes accelerate iron corrosion. Tools for genetic manipulation have only been developed for a few Fe(III)-reducing and methanogenic microorganisms known to corrode iron and in each case those microbes were found to accept electrons from Fe^0^ via direct electron transfer. However, iron corrosion is often most intense in the presence of sulfate-reducing microbes. The finding that *Desulfovibrio vulgaris* relies on H_2_ to shuttle electrons between Fe^0^ and cells revives the concept, developed in some of the earliest studies on microbial corrosion, that sulfate reducers consumption of H_2_ is a major microbial corrosion mechanism. The results further emphasize that direct Fe^0^-to-microbe electron transfer has yet to be rigorously demonstrated in sulfate-reducing microbes.

## Introduction

Microbial corrosion of iron-containing metals is a substantial economic problem, but the mechanisms are poorly understood (1–3). Sulfate reducers are often implicated in iron corrosion (1, 4–6). *Desulfovibrio vulgaris* has been the most studied pure culture isolate for investigating iron corrosion under sulfate-reducing conditions (7), dating back to the some of the earliest studies on microbial corrosion (8). Yet there is still substantial debate over how *D. vulgaris* corrodes iron. At least six mechanisms have been proposed (Figure 1).

**Figure 1.**
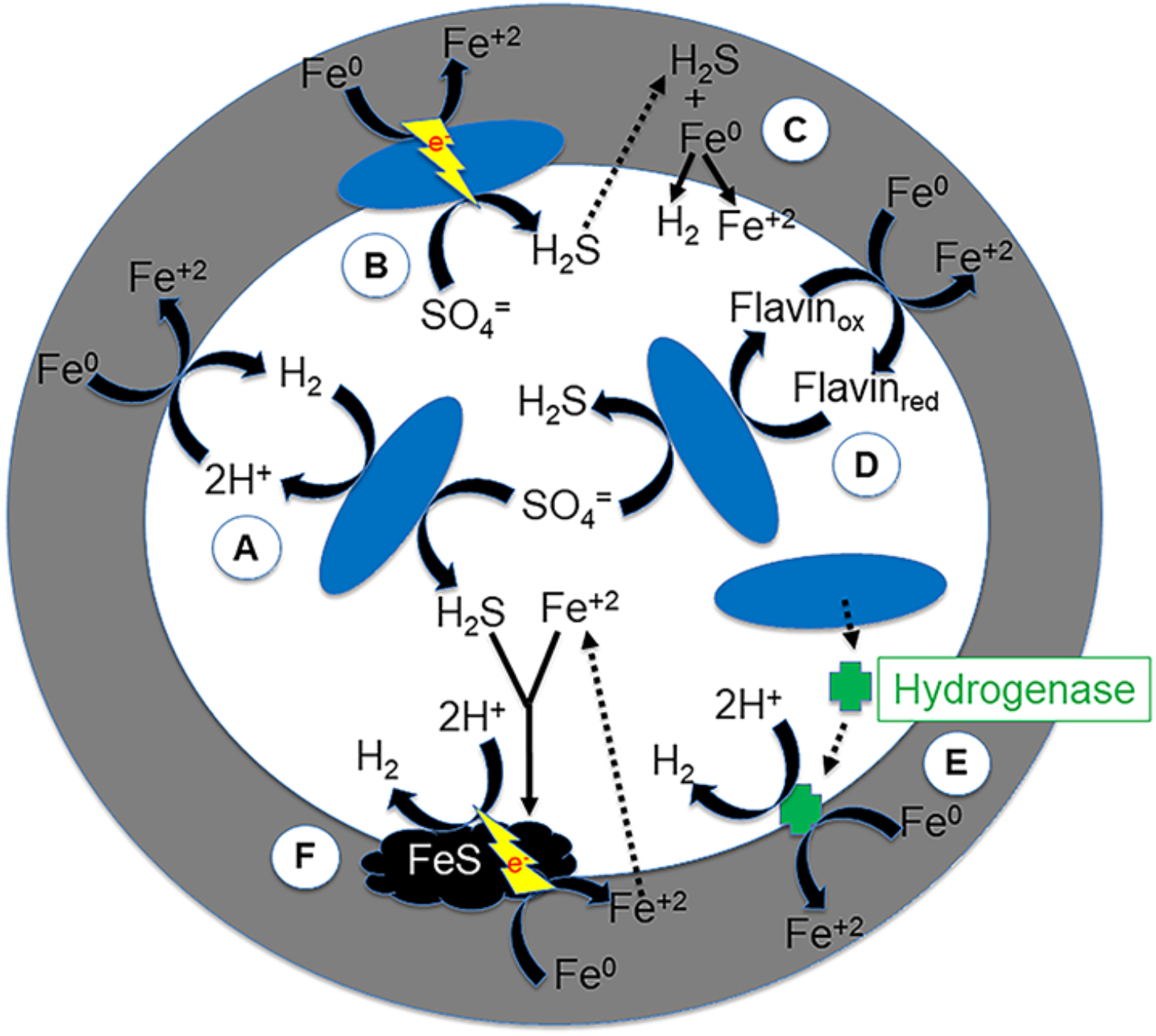
Previously proposed mechanisms for *Desulfovibrio vulgaris* to promote the oxidation of Fe^0^. (A) Consuming H_2_ produced from Fe^0^ abiotically reducing H^+^. (B) Direct Fe^0^-to-microbe electron transfer. (C) H_2_S produced from sulfate reduction reacting with Fe^0^ to produce H_2_. (D) Flavin serving as an electron shuttle to promote electron transfer from Fe^0^ to cells. (E) Hydrogenases released from cells catalyzing Fe^0^ oxidation via reduction of H^+^ to H_2_. (F) Iron sulfide functioning as a conductive conduit for improved electron transfer to H^+^ for abiotic H_2_ production.

The first mechanism proposed (8) was abiotic oxidation of Fe^0^ coupled to proton reduction to generate H_2_:

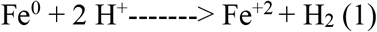

and consumption of the H_2_ produced via sulfate reduction:

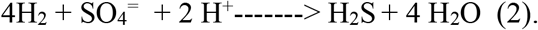

The H_2_ uptake might stimulate H_2_ production by lowering H_2_ concentrations, thus making H_2_ generation more thermodynamically favorable.

In a second mechanism, hydrogenases released from moribund cells might accelerate reaction #1 by catalyzing H_2_ production from Fe^0^ (9). A third proposed mechanism is that the H_2_S generated from *D. vulgaris* sulfate reduction might react with Fe^0^ to generate H_2_ (1):

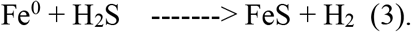

Alternatively Fe^+2^ and H_2_S can form iron sulfide precipitates that facilitate electron transfer from the Fe^0^ to H^+^, accelerating reaction #1 (2, 10).

Faster rates of corrosion following the addition of riboflavin (11–13) led to the suggestion that riboflavin can function as an electron shuttle that Fe^0^ reduces:

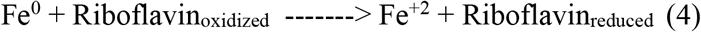

and *D. vulgaris* oxidizes the reduced riboflavin with the reduction of sulfate:

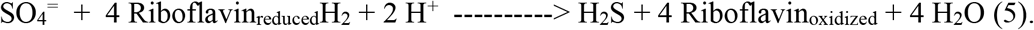

However, those studies did not determine whether Fe^0^ could donate electrons to riboflavin or whether reduced riboflavin can serve as an electron donor for sulfate reduction. The alternative possibility that riboflavin might stimulate other aspects of microbial metabolism was also not evaluated.

Direct Fe^0^-to-microbe electron transfer for *D. vulgaris* has also been proposed (14). Studies with *Geobacter* (15, 16), *Shewanella* (17, 18), and *Methanosarcina* (19) species have provided evidence for direct electron uptake from Fe^0^ by: 1) eliminating the possibility that H_2_ was serving as an electron shuttle between Fe^0^ and cells; and 2) demonstrating with gene deletions that outer-surface *c*-type cytochromes were required for electron uptake from Fe^0^. In contrast, no studies have previously reported on *D. vulgaris* corrosion with strains that were unable to use H_2_ (7). Furthermore, *D. vulgaris* lacks outer-surface cytochromes (20) and no other *D. vulgaris* outer surface electrical contacts are known. Unlike the microbes previously shown to directly accept electrons from Fe^0^ (15–19), *D. vulgaris* does not directly reduce Fe(III) (21), a capability common to most electroactive microbes (22).

The suggestion that *D. vulgaris* might be capable of direct Fe^0^-to-microbe electron transfer is related to an earlier suggestion that *D. ferrophilus* has this capability (23). However, direct Fe^0^-to-microbe electron transfer by *D. ferrophilus* was only inferred (23). *D. ferrophilus* uses H_2_ as an electron donor and the possibility of H_2_ serving as intermediary electron carrier between Fe^0^ and *D. ferrophilus* was not rigorously ruled out those early studies (3). In subsequent studies, *D. ferrophilus* grew with pure Fe^0^ as the electron donor, but not with stainless steel (24). This distinction is important because pure Fe^0^ abiotically generates H_2_ via reaction #1 (15, 25), but stainless steel does not (16). In contrast to *D. ferrophilus*, stainless steel was an effective electron donor for *Geobacter* and *Methanosarcina* strains capable of direct electron uptake from Fe^0^ (16, 19, 24). Notably, protease digestion of *D. ferrophilus* extracellular proteins did not affect sulfate reduction rates with Fe^0^ as the electron donor (26), a result inconsistent with a microbe making direct electrical contact with Fe^0^. Therefore, the evidence available to date suggests that *D. ferrophilus* is most likely to accept electrons from Fe^0^ via a H_2_ intermediate (24).

A rigorous strategy to evaluate the possibility of H_2_ serving as an intermediary electron carrier is to determine whether strains unable to use H_2_ can respire with Fe^0^ as the sole electron donor (15, 16, 18, 19, 27). In instances in which the wild-type strain of interest can consume H_2_, this can be accomplished by deleting genes necessary for H_2_ uptake (15, 18, 27). A strain of *D. vulgaris* in which genes for all of the annotated hydrogenases on the genome were deleted is available as one of a large collection of mutant strains (28). We report here on studies on Fe^0^-dependent sulfate reduction conducted with this hydrogenase-deficient strain.

## Results and Discussion

### Hydrogenase mutant unable to grow with H_2_ as electron donor

The hydrogenase mutant grew as well as the parental strain in medium with lactate as the electron donor and sulfate as the electron acceptor (Figure 2A), but unlike the parental strain, the hydrogenase mutant did not grow in medium with H_2_ as the sole electron donor (Figure 2B). These results suggested that the hydrogenase mutant was a suitable strain to evaluate the role of H_2_ as an intermediary electron carrier during growth with Fe^0^ as the electron donor.

**Figure 2.**
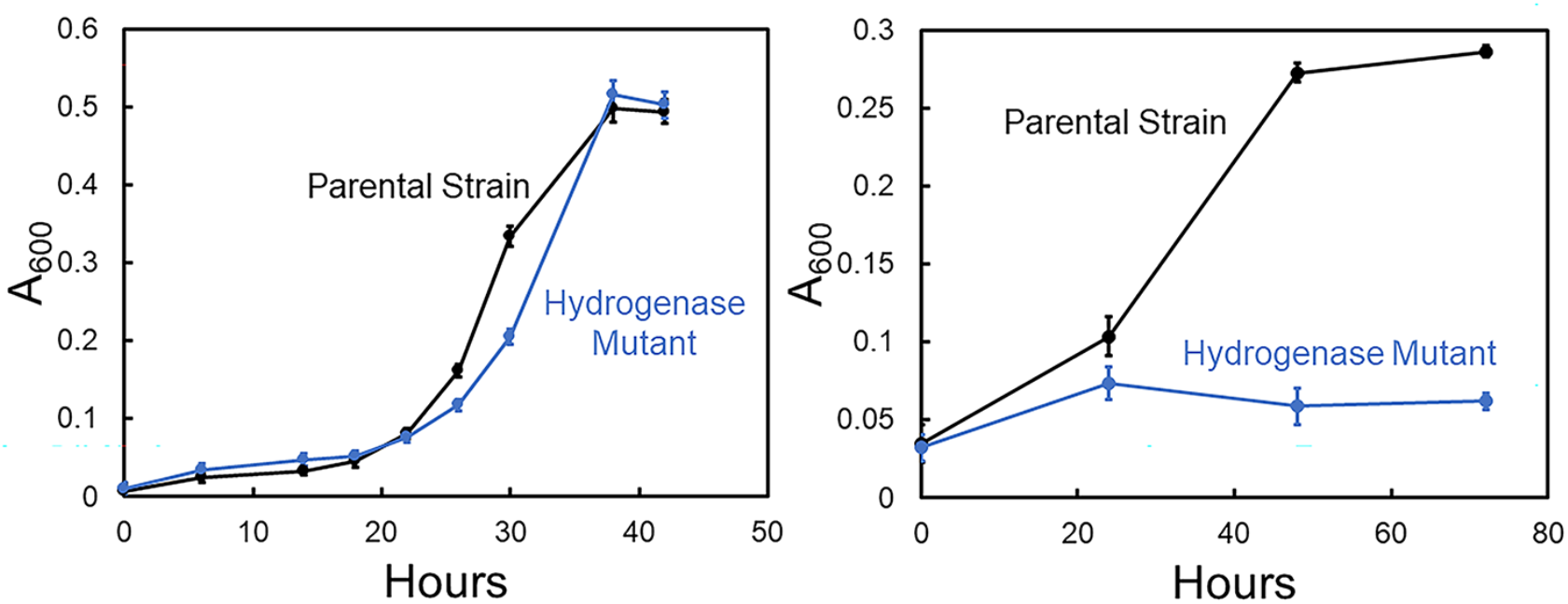
Growth of *Desulfovibrio vulgaris* parental strain and hydrogenase mutant with sulfate as the electron acceptor and either lactate (A) or H_2_ (B) as the electron donor, as measured by culture turbidity. Data are means and standard deviations of triplicate incubations.

### Hydrogenase mutant cannot reduce sulfate with Fe^0^ as electron donor

The parental strain reduced sulfate with Fe^0^ as the sole electron donor, but the hydrogenase mutant did not (Figure 3A). The slight decline in sulfate over time in cultures with the hydrogenase mutant could be attributed to carry over of lactate with the inoculum because the final sulfate levels for the hydrogenase mutant with Fe^0^ were the same as for the parental strain without Fe^0^ (Figure 3A). As expected from previous studies under similar conditions (15), H_2_ accumulated in sterile controls (Figure 3B) reflecting abiotic Fe^0^ oxidation coupled to H^+^reduction. H_2_ also accumulated in cultures inoculated with the hydrogenase mutant, further demonstrating the inability of this strain to consume H_2_. H_2_ accumulated more in the hydrogenase mutant cultures than in the uninoculated control, probably due to the sulfide that was transferred along with the inoculum (see sulfide effect on H_2_ production below). In contrast, the parental strain maintained low H_2_ concentrations (Figure 3B), as expected for a microbe that can consume H_2_ produced from Fe^0^ (15). In the presence of Fe^0^, sulfate reduction of the parental strain declined (Figure 3A) as H_2_ production plateaued in abiotic controls (Figure 3B), consistent with H_2_ serving as the electron donor for sulfate reduction. These results indicated that H_2_ produced from Fe^0^ was an important electron donor for sulfate reduction by the parental strain.

**Figure 3.**
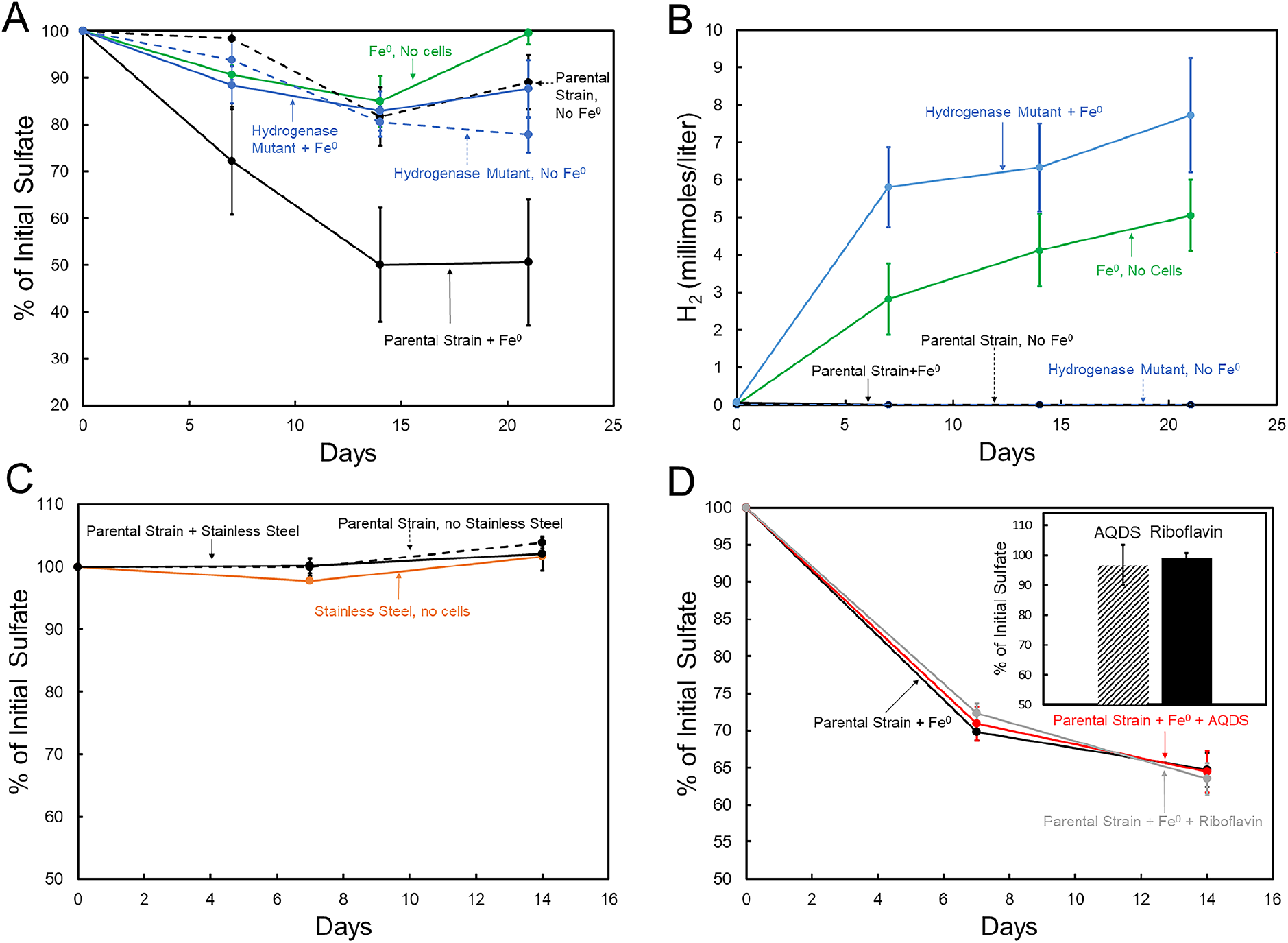
Sulfate reduction and H_2_ concentrations with iron as the electron donor in the presence of *Desulfovibrio vulgaris* parental strain or hydrogenase mutant. Sulfate loss (A) or H_2_ concentrations (B) over time with pure Fe^0^ as the sole electron donor in the presence of the *D. vulgaris* strains and in controls without cells or without Fe^0^. (C) Lack of sulfate depletion with 316L stainless steel as the potential electron donor for the parental strain and in controls without cells or without stainless steel. (D) Sulfate loss over time in parental strain cultures amended with riboflavin, anthraquinone-2,6-disulfonate (AQDS), or with no amendments, with Fe^0^ as the sole electron donor. The bar graph inset in (D) shows lack of sulfate loss in cultures of the hydrogenase mutant with Fe^0^ as the electron donor and amendments of riboflavin or AQDS. Data are means and standard deviations of triplicate incubations.

However, the quantity of H_2_ that accumulated in abiotic Fe^0^-only controls or in the presence of Fe^0^ and the hydrogenase mutant was not sufficient to account for the amount of sulfate that the parental strain reduced with Fe^0^ as the electron donor. For example, on day 14 the parental strain had reduced 3.4 mM sulfate (50% of the time 0 concentration of 6.8 mM), which would require 13.6 mM H_2_ (4:1 stoichiometry of H_2_ oxidized per sulfate reduced, reaction #2). Only ca. half that much H_2_ accumulated in the hydrogenase mutant cultures (Figure 3B). One possibility for this disparity is that because rapid H_2_ uptake by the parental strain maintained low H_2_ concentrations (Figure 3B), H_2_ production from Fe^0^ (reaction #1) was more thermodynamically favorable, possibly accelerating H_2_ generation over that in the hydrogenase mutant cultures in which H_2_ accumulated.

### Sulfide stimulates H_2_ production from Fe^0^

Sulfide that the parental strain generated from sulfate reduction with Fe^0^ as the electron donor is also likely to have promoted H_2_ production (Figure 4). Parental strain sulfide production was evident from the intense black precipitates indicative of iron sulfides on the Fe^0^ (Figure 4A). In contrast, there was only a small amount of iron sulfide on the Fe^0^ of the hydrogenase mutant cultures, which could be attributed to sulfide transferred along with the inoculum (Figure 4A). Sulfide was added to sterile medium, generating black iron sulfide precipitates (Figure 4B), to assess the possible sulfide impact on H_2_ production. Adding sulfide stimulated H_2_ generation (Figure 4C). One potential source of more H_2_ was the reaction of sulfide with Fe^0^ (reaction #3) in which there is a 1:1 stoichiometry for sulfide reacted and H_2_ produced. However, within 300 h the addition of 1.25 mM sulfide produced 2.7 mmol/liter H_2_ (Figure 4c), more than twice that expected from reaction #3. This result suggested that, as previously proposed (2, 10), iron sulfide precipitates also facilitated electron transfer from Fe^0^ to H^+^ (reaction #1), leading to additional H_2_ formation. Addition of 10-fold more sulfide only increased H_2_ an additional ca. 2-fold (Figure 4C), further demonstrating a lack of defined stoichiometry between sulfide additions and H_2_ formation.

**Figure 4.**
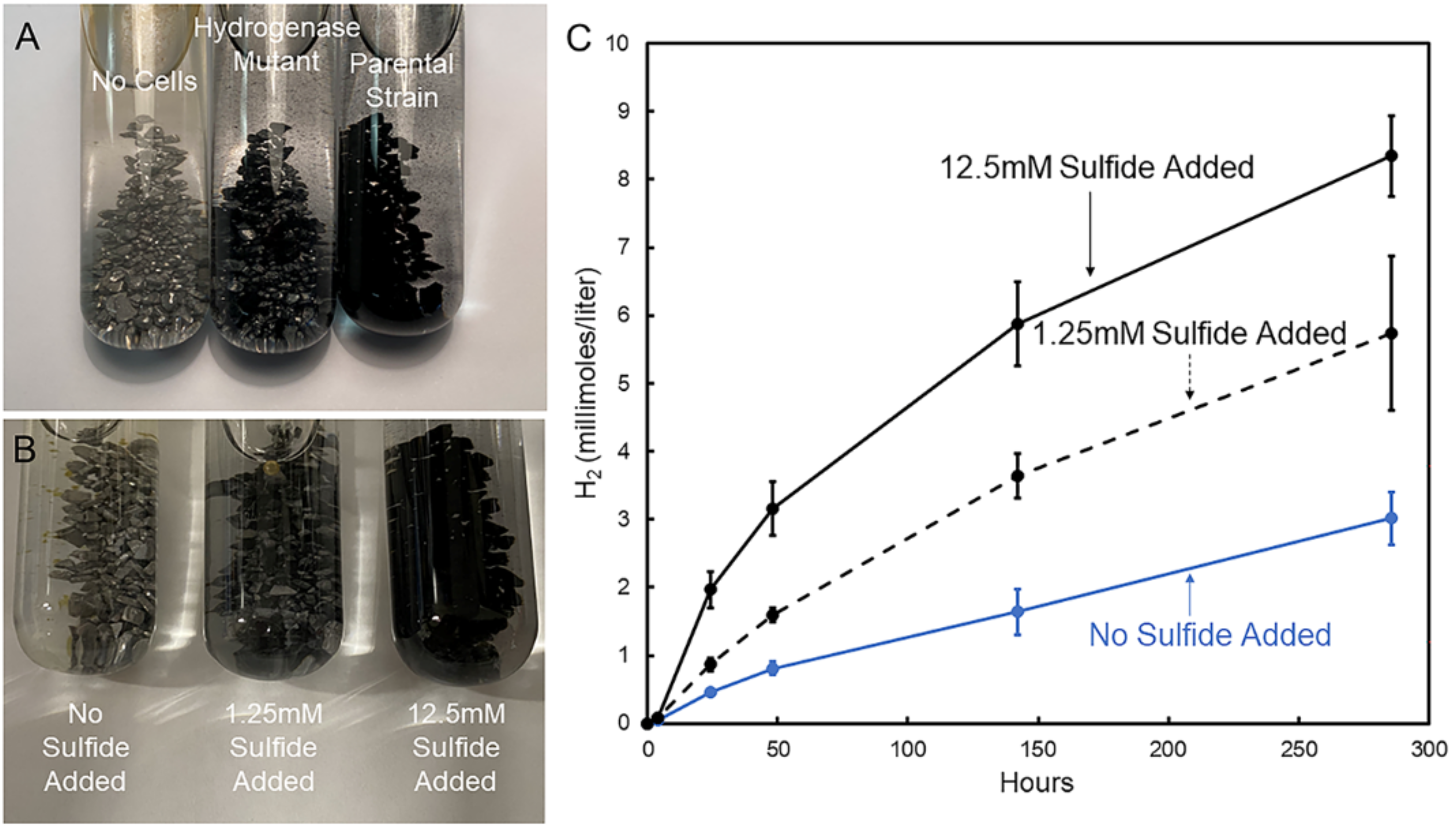
Iron sulfide accumulations in culture and impact of sulfide on abiotic H_2_ production. (A) Appearance of uninoculated Fe^0^-containing medium and cultures inoculated with either the hydrogenase mutant or parental strain after 14 days of incubation. (B) Appearance of sterile Fe^0^-containing medium amended with 1.25 or 12.5 mM final concentration of sodium sulfide. (C) Accumulation of H_2_ over time in sterile media with and without added sulfide. Data are means and standard deviations of quadruplicate incubations.

### Culture supernatant does not stimulate H_2_ production

Hydrogenases released from some microbes can accelerate H^+^ reduction with Fe^0^ (27, 29, 30) and hydrogenase activity has been detected in supernatants of moribund *D. vulgaris* cultures (9). However, supernatants from *D. vulgaris* cultures grown either with H_2_ or Fe^0^ did not stimulate H_2_ production from Fe^0^ over that in abiotic controls.

### Stainless steel studies further indicate the importance of H_2_ as electron carrier

The inability of the hydrogenase mutant to reduce sulfate with Fe^0^ as the electron donor contrasts with electroactive microbes such as *Geobacter sulfurreducens* (15) or *Shewanella oneidensis* (18), which continue to utilize Fe^0^ as an electron donor even after gene deletions have eliminated the capability for H_2_ uptake. Both *G. sulfurreducens* and *S. oneidensis* are capable of direct electron uptake as evidenced from an inhibition of Fe^0^-based respiration when genes for key outer-surface *c*-type cytochromes are deleted (15, 17, 18). Thus, the lack of sulfate reduction by the *D. vulgaris* hydrogenase mutant suggests that it is unlikely to support sulfate reduction with direct electron uptake from Fe^0^.

This conclusion was further supported with the results of studies in which stainless steel was provided as the electron donor. Unlike pure Fe^0^, H_2_ production from stainless steel is minimal (16). However, microbes capable of direct electron uptake from Fe^0^ can extract electrons from stainless steel to support anaerobic respiration (16, 17, 19). *D. vulgaris* did not reduce sulfate with stainless steel as the electron donor (Figure 3C).

### Electron shuttles do not promote Fe^0^-dependent sulfate reduction

An alternative proposed electron transfer mechanism in Fe^0^ corrosion is that flavins shuttle electrons between Fe^0^ and *D. vulgaris* (Figure 1). An observed increase in Fe^0^ corrosion when riboflavin is added to *D. vulgaris* cultures has been offered as evidence for flavin shuttling (11–13). However, the riboflavin amendments were to complex medium in which lactate was provided as an electron donor in addition to Fe^0^. It was not demonstrated that the riboflavin additions increased rates of Fe^0^-dependent sulfate reduction. In order to examine the possibility of electron shuttles facilitating electron transfer between Fe^0^ and *D. vulgaris*, studies were conducted under defined conditions with Fe^0^ as the sole electron donor for sulfate reduction and either riboflavin or the known electron shuttle anthraquinone-2,6,-disulfonate (AQDS) (31, 32). Riboflavin or AQDS did not accelerate Fe^0^-dependent sulfate reduction in the parental strain and did not enable the hydrogenase mutant to reduce sulfate with Fe^0^ as the electron donor (Figure 3D). The mid-point potentials of AQDS (−184 mV) and riboflavin (−208 mV) are probably too positive for the reduced form of these molecules to support the reduction of sulfate to sulfide (midpoint potential −217 mV). Therefore, the enhanced *D. vulgaris* Fe^0^ corrosion with riboflavin amendments (11–13) is likely to represent an impact of riboflavin on some aspect of growth or metabolism other than enhancement of electron transfer from Fe^0^ via an electron shuttle.

### *D. vulgaris* attaches to Fe^0^ electron donor

The turbidity of *D. vulgaris* growing on Fe^0^ was very low compared to the turbidity in H_2_-grown cultures when a comparable amount of sulfate had been reduced (Figure 5A). Confocal scanning laser microscopy revealed that cells attached to the Fe^0^ surface (Figure 5B,C). The attachment of cells places cells at the point of H_2_ production, which should be advantageous because it enables H_2_ uptake at the point of production where localized H_2_ concentrations are higher than in the bulk surrounding environment. Furthermore, localized conditions at the cell/Fe^0^ interface are likely to accelerate Fe^0^ oxidation (Figure 5D). For example, attached *D. vulgaris* oxidizing H_2_ can make Fe^0^ oxidation more thermodynamically favorable, both by removing a product of the reaction (H_2_) and resupplying a reactant (H^+^) near the Fe^0^ surface. Sulfide produced at the Fe^0^ surface can further accelerate H_2_ production.

**Figure 5.**
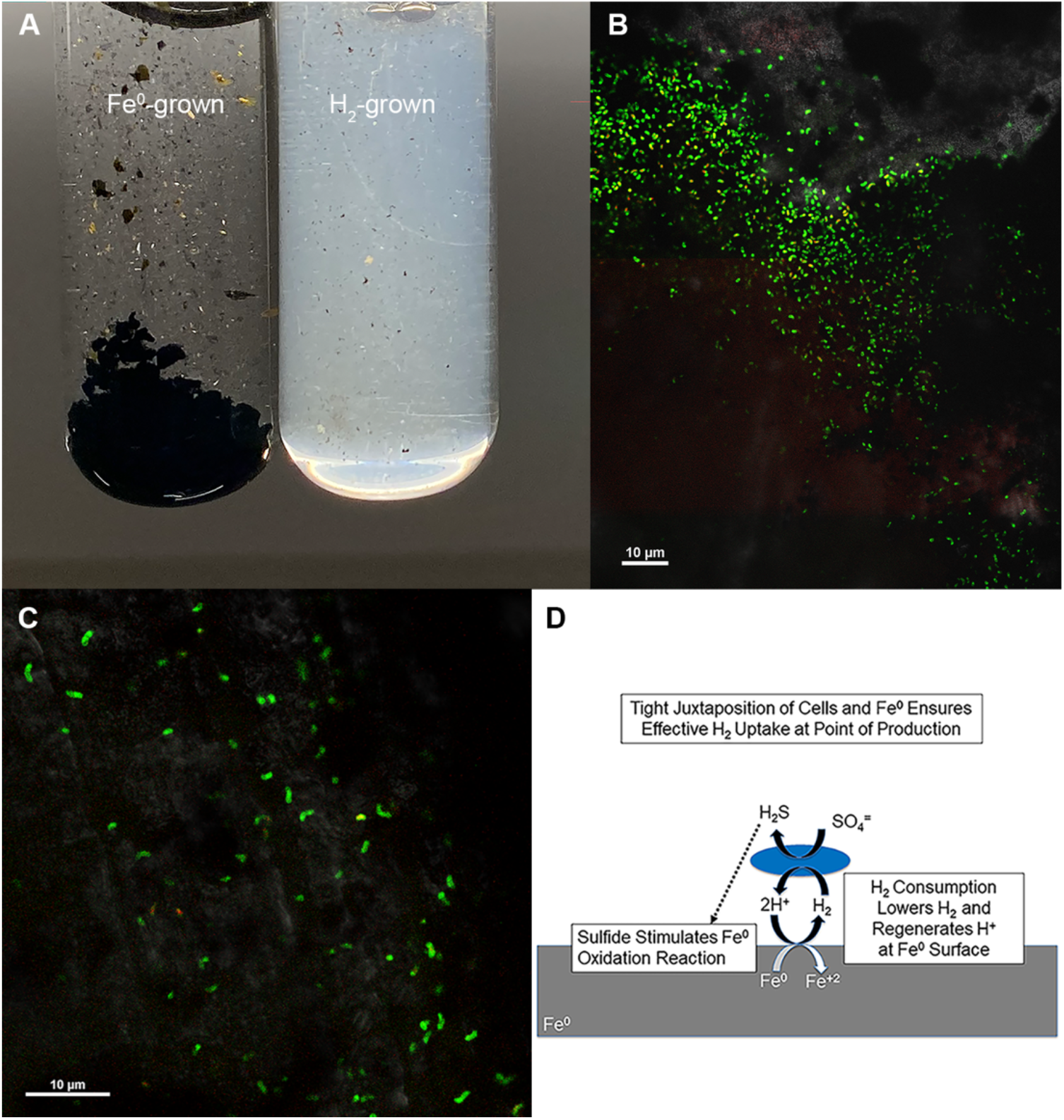
*Desulfovibrio vulgaris* attached to Fe^0^ serving as sole electron donor. (A) Lack of cell turbidity of culture growing with Fe^0^ as the source of H_2_ versus culture turbidity in culture grown under an atmosphere of H_2_. (B,C) Confocal scanning laser microscopy images showing cells attached to the Fe^0^ surface. (D) Mechanisms by which attachment to Fe^0^ might accelerate H_2_ production.

A common practice in Fe^0^ corrosion studies has been to infer that corrosion rates faster than that observed from abiotic H_2_ generation are indicative of corrosion mechanisms other than H_2_ serving as an intermediary electron carrier between Fe^0^ and cells (3). However, the possibilities for attached H_2_-consuming cells to accelerate H_2_ production from Fe^0^ illustrate the limitations to that reasoning.

### Implications

Understanding how *D. vulgaris* promotes Fe^0^ oxidation is important because it is the microbe that has been used to develop much of the existing mechanistic framework to describe how sulfate reducers corrode Fe^0^ (7). The results demonstrate that the primary mechanism for *D. vulgaris* to reduce sulfate with Fe^0^ as an electron donor is with H_2_ serving as an electron shuttle between Fe^0^ and the cells. Sulfate was not reduced in the absence of genes required for H_2_ uptake, even when previously proposed organic electron shuttles were added. All the microbes that have been previously shown to be capable of direct electron uptake from Fe^0^ have outer-surface *c*-type cytochromes known to be involved in extracellular electron exchange with other donors/acceptors (15–19). *D. vulgaris* lacks outer-surface *c*-type cytochromes (20). Direct electron uptake from extracellular electron donors by routes other than cytochromes is possible (33). For example, several methanogen species that lack outer-surface *c*-type cytochromes appear to directly accept electrons from *Geobacter metallireducens* (34–37). However, the results presented here demonstrate that *D. vulgaris* does not function as an electrotroph with Fe^0^ as the electron donor. If *D. vulgaris* is representative of the sulfate reducers most responsible for the corrosion of ferrous metals, then potent hydrogenase inhibitors might provide a targeted approach to mitigate iron corrosion.

However, microbes other than sulfate reducers are also likely to contribute to corrosion (3, 30, 38, 39). Elucidating the mechanisms by which a diversity of microbes accelerate corrosion is essential for understanding why corrosion takes place, predicting corrosion rates under various environmental conditions, and developing strategies for corrosion prevention. The studies reported here further demonstrate that construction of appropriate mutants is a powerful approach to distinguish between a complexity of potential corrosion mechanisms.

## Methods and Materials

### Microbial Strains

*Desulfovibrio vulgaris* strains JW710 and JW5095, which were constructed in the laboratory of Judy Wall, University of Minnesota (28, 40), were provided from a repository of *D. vulgaris* mutants by Valentine V. Trotter and Adam M. Deutschbauer of the Lawrence Berkeley Laboratory. Strain JW710, is as a platform strain for a markerless genetic exchange system in *D. vulgaris* (40). The *upp* gene encoding uracil phosphoribosyltransferase has been deleted, to enable utilization of the *upp* gene as a counterselectable marker (40). Strain JW5095 was constructed by markerless deletion of all the hydrogenases that have been described in the *D. vulgaris* genome: DVU1921-22, DVU2525-26, DVU1917-18, DVU1769-70, DVU0429-34, DVU2286-93, and DVU1771 (28).

### Culture Conditions

Cultures were routinely grown anaerobically at 37 °C in 10 ml of medium in 28 ml anaerobic pressure tubes (Bellco, Inc.), under N_2_/CO_2_ (80:20) in a modification of the previously described NBAF medium (41), designated NB medium. Per liter of deionized water NB medium contains: 0.42 g of KH_2_PO_4_, 0.22 g of K_2_HPO_4_, 0.2 g of NH_4_Cl, 0.38 g of KCl, 0.36 g of NaCl, 0.04 g of CaCl_2_ · 2H_2_O, 0.1 g of MgSO_4_ · 7H_2_O, 1.8 g of NaHCO_3_, 0.5 g of Na_2_CO_3_,, 1.0 ml of 1 mM Na_2_SeO_4_, 15.0 ml of a vitamin solution (42), and 10.0 ml of NB trace mineral solution. The composition of the NB trace mineral solution per liter of deionized water is 2.14 g of nitriloacetic acid, 0.1 g of MnCl_2_ · 4H_2_O, 0.3 g of FeSO_4_ · 7H_2_O, 0.17 g of CoCl_2_ · 6H_2_O, 0.2 g of ZnSO_4_ · 7H_2_O, 0.03 g of CuCl_2_ · 2H_2_O, 0.005 g of AlK(SO_4_)_2_ · 12H_2_O, 0.005 g of H_3_BO_3_, 0.09 g of Na_2_MoO_4_, 0.11 g of NiSO_4_ · 6H_2_O, and 0.02 g of Na_2_WO_4_ · 2H_2_O. Cells were routinely grown with sodium DL-lactate as the electron donor (20 mM) and sodium sulfate (20 mM) as the electron acceptor. Growth was monitored by inserting culture tubes directly into a spectrophotometer and determining A_600_. Growth with H_2_ as the sole electron donor was evaluated with 5 mM sodium acetate as a carbon source and H_2_ (140 kPa) as the sole electron donor. Cultures were routinely repressurized with H_2_ to compensate for any H_2_ consumption.

To evaluate growth with Fe^0^ as the potential electron donor, cells were grown in NB medium with Fe^0^ granules (2 g;1 to 2 mm diameter; Thermo Scientific) as the sole electron donor, 5 mM sulfate as the electron acceptor, and 5 mM sodium acetate as a carbon source. When specified, 50 μM riboflavin or 50 μM anthraquinone-2,6-disulfonate were added from concentrated anaerobic stock solutions. For studies with 316L stainless steel as the potential electron donor for sulfate reduction, 5 stainless steel cubes (5 mm x 3 mm x 3 mm) replaced the pure Fe^0^. The stainless steel cubes were polished with sand paper and the pure Fe^0^ and stainless steel were presterilized with ethanol as previously described (24).

### Impact of Added Sulfide or Culture Supernatant on H_2_ Production

A final concentration of either 1.25 mM or 12.5 mM sodium sulfide was added to sterile Fe^0^-containing medium to determine whether sulfide stimulated H_2_ production. Culture filtrates were prepared by filtering late log grown cultures (Fe^0^-grown or H_2_-grown) through a 0.2 μM PES filter in a Coy anaerobic glove bag (gas phase 7:20:73 H_2_/CO_2_/N_2_) into pressure tubes with 2 g of Fe^0^. Tubes were resealed and flushed with N_2_:CO_2_ (80:20) for 5 minutes. Controls were sterile NB medium.

### Analytical Methods

For sulfate determinations, culture aliquots (0.1 ml) were anaerobically withdrawn with a syringe and needle, filtered (0.22 μm, PVDF), and analyzed with a Dionex ICS-1000 with an AS22 column and AG22 guard with an eluent of 4.1 mM sodium carbonate and 1 mM sodium bicarbonate at 1.2 ml/min. H_2_ concentrations in the headspace were monitored on an Agilent 6890 gas chromatograph fitted with a thermal conductivity detector. The column was a Supelco Carboxen 1010 plot capillary column (30 m x 0.53 mm) with N_2_ carrier gas and 0.5 ml injections. The oven temperature was 40°C and the inlet was splitless at 5.5 psi and 225°C, the detector had a makeup flow of 7 ml/min and temperature of 225°C.

### Confocal Microscopy

For confocal microscopy, Fe^0^ was gently removed from the pressure tube, soaked in isotonic wash buffer for 10 minutes, drained, stained for 10 minutes (Live/Dead BacLight bacterial viability kit (Thermo Fisher) (1 ml staining with 3 μl of each stain per ml)), and destained for 10 minutes in isotonic wash buffer. Fe^0^ pieces were then mounted on petri plates with an antifade/glycerol mixture. Cells were visualized with a 100x objective on a Nikon A1R-SIMe confocal microscope with NIS-Elements software.

## Acknowledgements

These studies were made possible by the public availability *Desulfovibrio vulgaris* strains JW710 and JW5095 made in the laboratory of Judy Wall, University of Missouri. Thomas R. Juba constructed strain JW5095. We thank Valentine Trotter and Adam Deutschbauer of the Berkeley National Laboratory for providing us with strains used in this study. Confocal microscopy was performed in the Light Microscopy Facility and Nikon Center of Excellence at the Institute for Applied Life Sciences, UMass Amherst.

